# Population based hospitalization burden of laboratory-confirmed hand, foot and mouth disease caused by multiple enterovirus serotypes in southern China

**DOI:** 10.1101/403659

**Authors:** Shuanbao Yu, Qiaohong Liao, Yonghong Zhou, Shixiong Hu, Qi Chen, Kaiwei Luo, Zhenhua Chen, Li Luo, Wei Huang, Bingbing Dai, Min He, Fengfeng Liu, Qi Qiu, Lingshuang Ren, H. Rogier van Doorn, Hongjie Yu

## Abstract

**Background:** Hand, foot and mouth disease (HFMD) is spread widely across Asia, and the hospitalization burden is as yet not well understood. Here, we estimated serotype-specific and age-specific hospitalization rates of HFMD in Southern China.

**Methods:** We enrolled pediatric patients admitted to 3/3 county-level hospitals and 3/23 township level hospitals in Anhua county, Hunan (CN) with HFMD, and collected samples to identify enterovirus serotypes by RT-PCRs between October 2013 and September 2016. The information of other eligible but un-enrolled patients were retrospectively collected from the same six hospitals. Monthly number of hospitalizations for all causes was collected from each of 23 township level hospitals to extrapolate hospitalizations associated with HFMD among these.

**Results:** During the three years, an estimated 3,236 pediatric patients were hospitalized with lab-confirmed HFMD, and among these only one patient was severe. The mean hospitalization rates were 660 (95% CI: 638-684) per 100,000 person-years for lab-confirmed HFMD, with higher rates among CV-A16 and CV-A6 associated HFMD (213 vs 209 per 100,000 person-years), and lower among EV-A71, CV-A10 and other enteroviruses associated HFMD (134, 39 and 66 per 100,000 person-years, *p*<0.001). Children aged 12-23 months had the highest hospitalization rates (3,594/100,000 person-years), followed by those aged 24-35 months (1,828/100,000 person-years) and 6-11 months (1,572/100,000 person-years). Compared with other serotypes, CV-A6-associated hospitalizations were evident at younger ages.

**Conclusions:** Our study indicates a substantial hospitalization burden associated with non-severe HFMD in a rural county in southern China. Future mitigation policies should take into account the disease burden identified, and optimize interventions for HFMD.

## Introduction

Hand, foot and mouth disease (HFMD) is a common infectious disease that mainly affects children below 5 years of age [1]. HFMD is caused by multiple serotypes of Enterovirus species A, among which enterovirus 71 (EV-A71) and coxsackievirus A16 (CV-A16) are the most frequently detected [2]. EV-A71 is of particular concern as it can cause neurological and systematic complications, and even fatal outcomes [2–4]. Coxsackie virus A6 (CV-A6) and Coxsackie virus A10 (CV-A10) have become more prevalent among HFMD outbreaks. Their re-emergence was first identified in HFMD cases in Europe and Singapore between 2008 and 2011 [5–8]. They are also responsible for considerable cases of HFMD in China since 2013 [9, 10]. Severe complications associated with CV-A6 and CV-A10 have also been reported [11, 12]. Currently, no specific antiviral treatments are available for HFMD. Three inactivated monovalent EV-A71 vaccines have been licensed in mainland China, with high efficacy (90.0%-97.4%) against EV-A71-HFMD, but no cross-protection against other enterovirus serotype-associated HFMD [13, 14]. Bivalent and multivalent enterovirus vaccines are under development [15, 16].

Since 1997, HFMD has widely spread across Asia, including Malaysia, Japan, Singapore, Vietnam, Cambodia, and China [1]. Understanding the age and serotype-specific burden of HFMD, including the hospitalization burden, is valuable in informing healthcare systems, vaccine strategies and other intervention policies. However, the hospitalization burden of HFMD has not been thoroughly studied in a well-defined catchment population. Indirect estimates of hospitalization rates of HFMD are hampered by limited availability of population level incidence and unknown hospitalization rates among HFMD cases. Population level incidence of HFMD has been estimated in Japan [17], Singapore [18], Malaysia [19], and China [2, 20]. These estimates were based on notifiable surveillance data which may underestimate the true prevalence. The risk of hospitalization for HFMD varied between 1.3% and 24.3% [21–25], possibly due to patients with distinct severity and different causative serotypes of enterovirus in different studies. The number of people that were included in these studies ranged from 6,027 to 1,081,046 [21–25]. Additionally, there is an increasing threat of enterovirus serotypes of non-EV-A71 among both mild and severe HFMD [7, 11, 20]. Therefore, the hospitalization burden of HFMD caused by CV-A6, CV-A10, CV-A16 and other enterovirus serotypes requires further assessment.

We aim to estimate population level hospitalization rates of HFMD by age and enterovirus serotype in a well-defined catchment population in China between October 2013 and September 2016.

## Methods

### Study setting

This study was conducted in Anhua County, Yiyang Prefecture, Hunan Province, China [26]. The total population residing in Anhua County was 1,017,463 according to the 2015 census [27]. A total of 165,050 (16%) of the population were children aged <15 years, and of these 61,123 (6%) were aged <5 years [27]. There were three county-level hospitals and 23 township level hospitals in Anhua County where HFMD patients were admitted. According to vaccination records, the EV-A71 vaccine was initiated in Anhua County on July 10, 2016.

### Case definitions

A probable case of HFMD was defined as a patient with a rash on their hands, feet, limbs or buttocks, ulcers or vesicles in the mouth, with or without fever. A lab-confirmed case was defined as a probable case with laboratory evidence of enterovirus infection detected by real-time RT-PCR (reverse transcription-polymerase chain reaction) or nested RT-PCR. Hospitalized patients were defined as patients hospitalized during at least 24 hours of admission.

### Virological surveillance of HFMD

Virological surveillance of HFMD was conducted among six hospitals, including three county-level hospitals and three township level hospitals between October 2013 and September 2016. The six hospitals were selected as they admitted 87% of the reported HFMD patients from Anhua County during 2010-2012. The methods for the virological surveillance have been described elsewhere [26]. Briefly, the pediatric patients (aged 0-14 years) who were hospitalized for HFMD, were enrolled after parents/legal guardians provided verbal consent. Throat swabs and stool samples (or rectal swabs) were collected within 24 hours of enrollment. A standardized form was used to collect data, including basic demographic information, date of illness onset, types of samples (stool, throat swab, or rectal swab), complications, and clinical outcome.

Swabs were immediately placed in viral transport mediums, and all samples were stored at −70 °C until testing. Viral RNA was extracted using QIAamp Viral RNA Mini Kit (QIAGEN, Hilden, Germany). RNA from each sample was tested with generic primers and probes targeting pan-enterovirus, and specific primers and probes targeting EV-A71, CV-A16, and CV-A6. If a sample tested positive in the generic and negative in the specific RT-PCRs, nested RT-PCRs were used to amplify VP1 regions. If negative in amplifying VP1 regions, nested RT-PCRs on VP4-VP2 regions were used for further identification of enterovirus serotype.

### Estimation of hospitalization burden

Some patients who were admitted to hospital for HFMD during the study period were not captured by the above virological surveillance. Therefore, we retrospectively collected the monthly number of eligible but unenrolled hospitalized patients with diagnosis of HFMD by age group at each of the six hospitals. We extracted data from the electronic Hospital Information System (HIS) at the three county-level hospitals. We manually reviewed the paper-based medical charts at the three township hospitals. We did not visit the other 20 township hospitals to collect the number of HFMD-associated hospitalizations manually. Instead, the number of patients who were hospitalized between October 2014 and September 2016, the cause of their hospitalization and their age was extracted from the Rural Health Information System (RHIS) of Hunan Province (http://220.170.145.170:8888/chss/) for each of the 23 township hospitals.

A multiplier model [28] was used to estimate the hospitalization rates of HFMD in this study (Fig 1). We divided the patients and the population denominators into eight age groups, including <6 months, 6-11 months, 12-23 months, 24-35 months, 36-47 months, 48-59 months, 5-9 years, and 10-14 years. We assumed that the age stratified admission rates were comparable among the township level hospitals. We extrapolated the number of HFMD-associated hospitalizations per age group among 23 township level hospitals based on data from the three township hospitals and number of total hospitalizations among all township level hospitals (Fig 1). The total number of hospitalizations for HFMD in each age group in Anhua county was the sum of the number of hospitalizations captured and non-captured in virological surveillance among the county level hospitals, and number of hospitalizations among all township level hospitals. We estimated the number of hospitalizations attributable to lab-confirmed HFMD, EV-A71, CV-A16, CV-A6, CV-A10 and other enteroviruses (non-EV-A71 & non-CV-A16 & non-CV-A6 & non-CV-A10) associated with HFMD in Anhua County. We assumed that the age-specific distribution of enterovirus serotype in the enrolled group was the same as that in the non-enrolled group (Fig 1).

**Fig 1.**
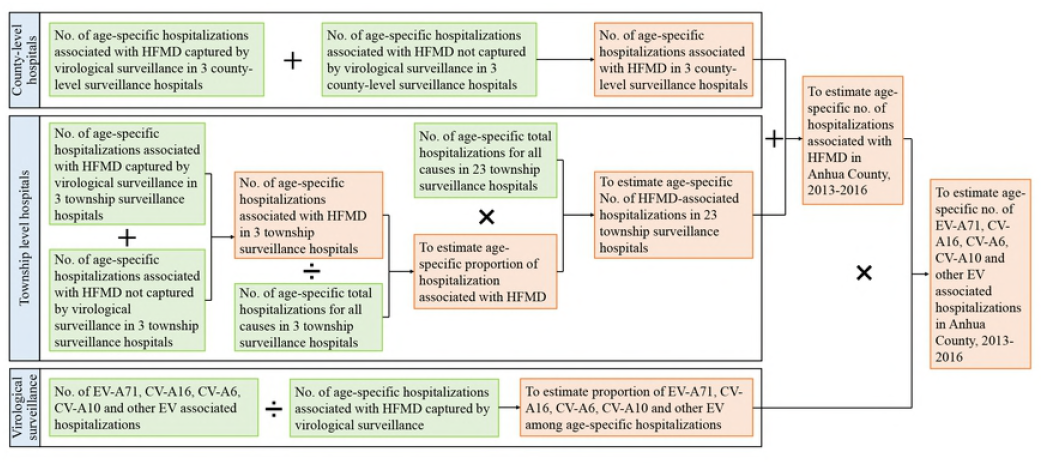
Flowchart of estimating age-specific and serotype-specific hospitalization burden associated with HFMD.

The hospitalization rates by serotype were estimated by dividing the serotype-specific number of hospitalizations by the size of the resident population. The age-specific population denominators of Anhua County from 2013 to 2016 were collected from the National Bureau of Statistics of China [27].

### Statistical analysis

The Poisson method was used to estimate 95% confidence intervals (CI) of hospitalization rates. The χ^2^ test or Fisher’s exact test was used to analyze categorical data. The Mann-Whitney U test was used to analyze ranked data. Student’s t test was used to analyze continuous data. Data cleaning and all analysis were conducted using R (version 3.4.2).

### Ethical approval

This study was approved by the Institutional Review Board (IRB) at the Chinese Center for Disease Control and Prevention (no. 201224). This study was considered to be part of a continuing public health outbreak investigation by National Health and Family Planning Commission of China and except from institutional review board assessment. Therefore, written informed consents were not obtained from subjects. The IRB approved the use of verbal consent, and agreed that we anonymized the specimens and personal information by permanently removing personal identifiers from the database. Anonymized samples were labeled with a random coding system. Verbal informed consent was obtained form the patients’ parents/guardians when sampling and filling out the questionnaire, and documented in forms.

## Results

### Virological surveillance

During the three-year virological surveillance, 2,836 (85%) of the total of 3,326 hospitalized patients with a diagnosis of HFMD were enrolled across the six hospitals. The baseline characteristics were similar between the enrolled and un-enrolled patients, including gender, age, and length of hospital stay [26]. One 27-month old child with detection of EV-A71 had symptoms of neurological involvement (frequent jittering and myoclonic jerks after 4 days of fever), and all others had uncomplicated illnesses. Enterovirus was detected among 2,517 (89%) patients via the real-time RT-PCR targeting pan-enterovirus. Nineteen serotypes of enterovirus were successfully identified in 2,513 (99.8%) patients. The most commonly detected were CV-A16 (33%, 819), CV-A6 (31%, 785), EV-A71 (20%, 514), and CV-A10 (6%, 149). CV-A6 infections were more frequently identified in patients younger than 2 years, while CV-A16 and EV-A71 accounted for a higher proportion of HFMD among children aged ≥3 years than CV-A6 (*p*<0.001) (S1 Table). A seasonal peak in total hospitalizations associated with HFMD was observed between April and June (S1 Fig).

### Serotype-specific hospitalization rates of HFMD

Between October 2013 and September 2016, an estimated 3,642 pediatric patients were hospitalized for HFMD in Anhua County. A total of 3,273 (90%) children were admitted in the three county-level hospitals and 369 (10%) children were admitted in the 23 township level hospitals (S2 Table). Among these, 3,236 (89%) patients were estimated to be positive for enterovirus. The mean hospitalization rates of probable HFMD were estimated as 743 (95% CI: 719-768) per 100,000 person-years, and the mean rates of lab-confirmed HFMD were 660 (95% CI: 638-684) hospitalizations per 100,000 person-years during the surveillance period (Fig 2A). Both the annual hospitalization rates of probable and lab-confirmed HFMD were highest during the 2013-2014 season (1,099 and 1,011 per 100,000 person-years), and lowest during the 2014-2015 season (458 and 422 per 100,000 person-years) (Fig 2B).

**Fig 2.**
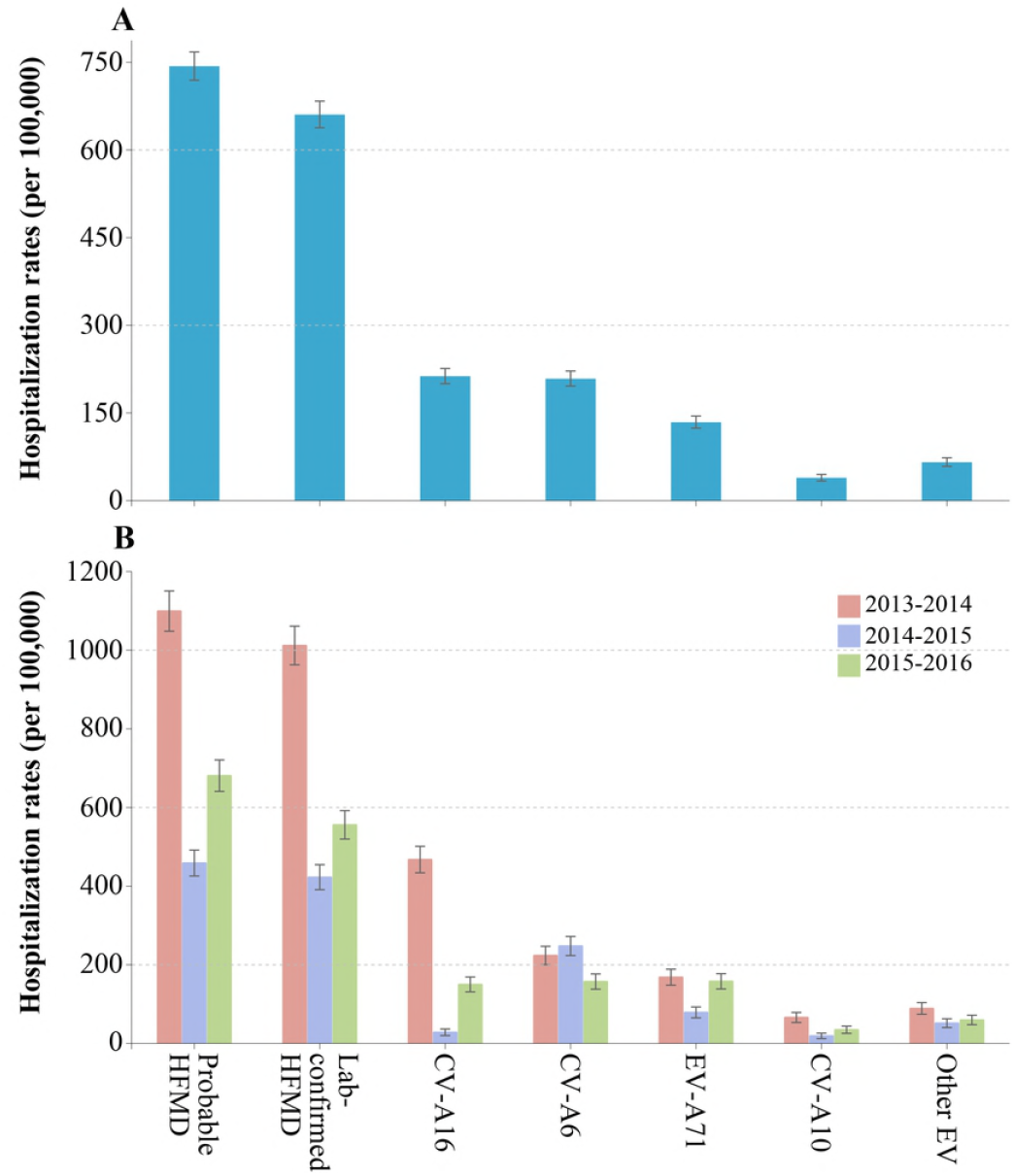
Hospitalization rates of HFMD in Anhua County, China, October 2013 - September 2016. (A) Average hospitalization rates of HFMD, overall and stratified by serotype. (B) Annual hospitalization rates of HFMD, overall and stratified by serotype.

Based on the assumption of similar distribution of serotypes among the enrolled and un-enrolled groups, 3,236 (89%) patients with lab-confirmed HFMD consisted of 1,043 with CV-A16, 1,023 with CV-A6, 657 with EV-A71, 191 with CV-A10, and 322 with other enteroviruses. The mean hospitalization rates were comparable between CV-A16-associated HFMD (213 per 100,000 person-years) and CV-A6-associated HFMD (209 per 100,000 person-years) during the surveillance period, and lower for EV-A71-associated HFMD (134 per 100,000 person-years), CV-A10-associated HFMD (39 per 100,000 person-years), and other enteroviruses (66 per 100,000 person-years) infections (*p*<0.001) (Fig 2A).

In the first year, between October 2013 and September 2014, the hospitalization rates for CV-A16 were the highest (467 per 100,000 person-years). While in the second year, CV-A6 had the largest burden of hospitalizations (248 per 100,000 person-years). The hospitalization rates were similar among CV-A16, CV-A6, and EV-A71 infections in the third year (*p*=0.568) (Fig 2B). The annual hospitalizations varied substantially over years for EV-A71, CV-A16, and CV-A10. However, CV-A6 displayed comparable hospitalization rates over three years (Fig 2B).

### Age-specific hospitalization rates of HFMD

The mean hospitalization rates of HFMD varied with age. The highest rates of lab-confirmed HFMD were among children aged 12-23 months (3,594 per 100,000 person-years), followed by 24-35 months (1,828 per 100,000 person-years) and then 6-11 months (1,572 per 100,000 person-years). The hospitalization rates were lower among infants younger than 6 months (381 per 100,000 person-years) and among children aged 5-14 years (84 per 100,000 person-years) (Fig 3B). Similar age distributions of hospitalization rates were observed for probable HFMD (Fig 3A). The mean rates of probable and lab-confirmed HFMD were 1,829 and 1,638 hospitalizations per 100,000 person-years (respectively) among children younger than 5 years. Unlike the distribution of yearly rates for children aged ≥12 months, the hospitalization rates decreased over three years for children aged 0-11 months (Fig 3C and D).

**Fig 3.**
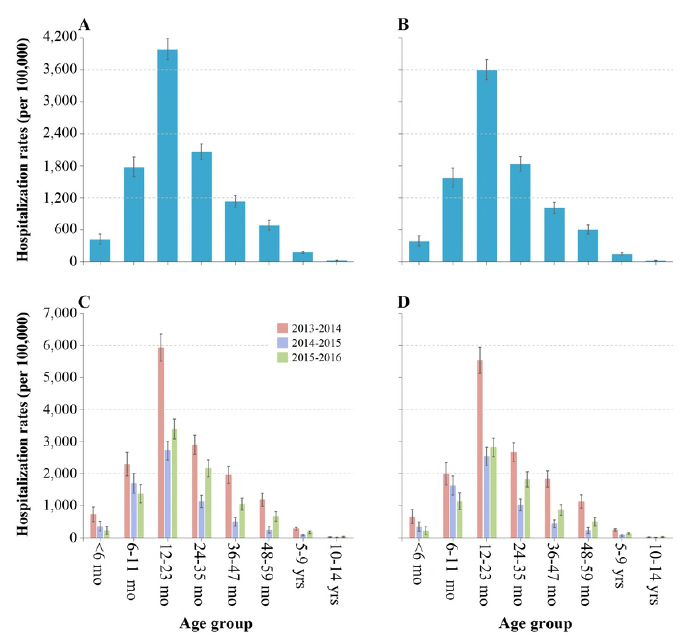
Age-specific hospitalization rates associated with HFMD in Anhua County, China, October 2013 - September 2016. (A) Average age-specific hospitalization rates of probable HFMD. (B) Average age-specific hospitalization rates of lab-confirmed HFMD. (C) Annual age-specific hospitalization rates of probable HFMD. (D) Annual age-specific hospitalization rates of lab-confirmed HFMD.

The four common enterovirus serotypes had highest hospitalization rates among children aged 12-23 months (955 for CV-A16, 1,344 for CV-A6, 640 for EV-A71, and 267 for CV-A10 per 100,000 person-years) (Fig 4A-D). CV-A6 uniquely had higher rates among children aged 6-11 months than among those aged 24-35 months (*p*<0.001), while CV-A10 had comparable rates between these age groups (*p*=0.849) (Fig 4A-D). Statistical analyses suggested that HFMD hospitalization associated with CV-A6 were evident at younger ages, compared to EV-A71 and CV-A16. The distribution pattern of age-specific hospitalization rates for CV-A16, EV-A71, and CV-A10 were consistent across the three years, between October 2013 to September 2016 (S2 Fig). For CV-A6, the higher hospitalization rates in 6-11 months than 24-35 months was not evident in the third year, between October 2015 to September 2016 (*p*=0.777) (S2 Fig).

**Fig 4.**
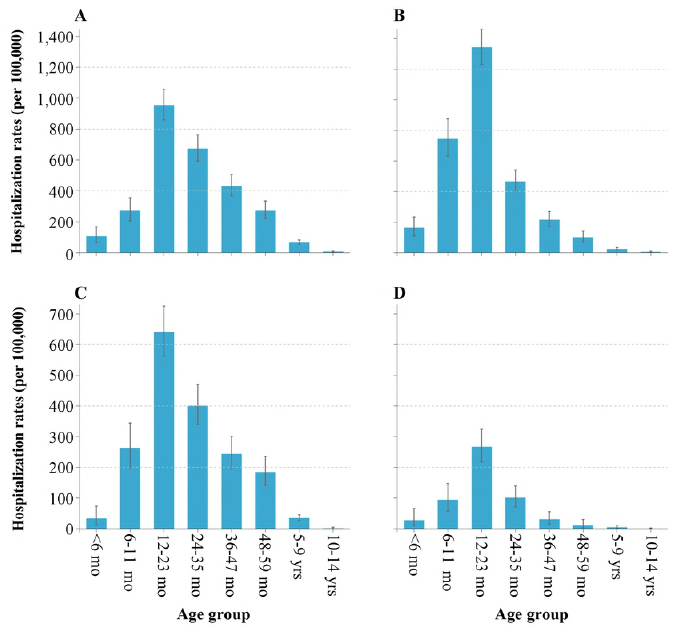
Age-specific and serotype-specific hospitalization rates associated with HFMD in Anhua County, China, October 2013 - September 2016. (A) Average age-specific hospitalization rates of CV-A16-associated HFMD. (B) Average age-specific hospitalization rates of CV-A6-associated HFMD. (C) Average age-specific hospitalization rates of EV-A71-associated HFMD. (D) Average age-specific hospitalization rates of CV-A10-associated HFMD.

## Discussion

### Principal findings

This study provides a comprehensive estimate of the hospitalization rates of probable and lab-confirmed HFMD in Anhua County, Hunan Province, China, between October 2013 and September 2016. An average of 743 probable HFMD and 660 lab-confirmed HFMD associated hospitalizations were estimated per 100,000 person-years in Anhua County, with the highest annual rates during 2013-2014 and the lowest during 2014-2015. CV-A16 and CV-A6 were associated with most (64%) lab-confirmed HFMD hospitalizations, and had higher hospitalization rates than EV-A71, CV-A10 and other enteroviruses during the study period. Hospitalization rates peaked among children aged 12-23 months, and decreased with age among EV-A71, CV-A16, CV-A6, CV-A10 and other enterovirus associated HFMD. Compared to other non-CV-A6 serotypes, CV-A6 had higher hospitalization rates in children aged 6-11 months than in children aged 24-35 months.

### Strength and comparison with previous studies

Because of the high proportion of asymptomatic enterovirus infections (EV-A71, 64.1-71.4%) in children [29, 30], and the relative scarcity (1.1%) of HFMD associated severe illness [2, 29], the burden of HFMD is substantially underestimated if symptomatic case notifications are only used. In this study, we actively captured HFMD-associated hospitalizations in all healthcare facilities in Anhua county instead of relying on passive surveillance. Additionally, our virological surveillance captured 78% of the total HFMD-associated hospitalizations over three years. This allowed us to attain an accurate representation of the enterovirus serotypes that cause HFMD. Samples were collected in a timely fashion; multiple samples were available and intensive methods were used to identify enterovirus serotypes. This enabled us to make a robust estimation of the population-based hospitalization burden stratified by age group and enterovirus serotype.

In our study, all children with mild illness (except one) were admitted to hospital. The reasons underlying overuse of healthcare resources for mild illnesses include a low hospitalization threshold applied by local physicians to capture all possible sudden deteriorations [3], parents self-requesting hospital admission and the rural healthcare insurance (New Rural Cooperative Medical Scheme) providing higher reimbursement for inpatient than outpatient care [31]. Proportions of mild patients accounted for 80-99% among HFMD-associated hospitalizations in previous reports [21, 24, 25, 29]. The patients without complications were hospitalized in these reports because of high fever, poor feeding, mouth ulcers, vomiting, or dehydration [24]. This indicates that hospitalizations for non-severe HFMD is common in China.

To our knowledge, population-based hospitalization rates for HFMD have not been reported until now. As the threshold for hospitalization of HFMD in our study was relatively low, we made comparisons with the incidence rates estimated using the notifiable surveillance data [2, 17–20]. The hospitalization rates estimated in our study (119/100,000 person-years) were comparable to the incidence rates reported in mainland China during 2008-2015 (127/100,000 person-years) [2, 20]. The rates we estimated were higher than the incidence rates in Malaysia during 2011-2014 (20-90/100,000 person-years) [19], and lower than the incidence rates in Singapore during 2001-2007 (126-436/100,000 person-years) [18], and Japan during 2002-2005 (743 vs 2,940-5,740 per 100,000 population) in children aged 0-14 years [17]. The variation in results could be associated with distinct activity intensity of enterovirus serotypes in different study periods, and differences among the surveillance systems. The dual layers of doctor-driven and teacher-driven surveillance make the data on notified symptomatic presentations of HFMD collected by the Singapore’s Ministry of Health unusually complete [29]. The surveillance of HFMD in China and Malaysia rely on all hospitals [2, 19]. The surveillance targeted HFMD in Japan are based on sentinel medical institutions according to the guidelines for surveillance of infectious diseases [17]. Outpatients were not included in our study, therefore the hospitalization rates we report could be an underestimate of the true incidence rates in Anhua County.

The age pattern of HFMD-associated hospitalizations was consistent with previous reports of HFMD-associated incidences [2, 19, 20]. Relatively low hospitalization rates among children younger than 6 months were likely due to protection by maternal antibodies [32]. Compared to older children who may attend kindergartens or nurseries, children aged 6-23 months have a lower intensity of contact with those of similar ages due to limited mobility. The high hospitalization rates among this age group may suggest higher susceptibility and/or possible transmission routes from contact with asymptomatic adults or contaminated environments [20]. Serological surveillance also revealed a low level of sero-prevalence in children aged 6-23 months [32, 33].

Our study found that CV-A6 presented a higher burden of HFMD-associated hospitalization than EV-A71. This was consistent with the increasing activity of CV-A6 in China [10, 11, 34–36] and the regional introduction of CV-A6 in 2012. CV-A6-associated hospitalizations were also higher among younger age groups than EV-A71 and CV-A16 [37–39]. This may result from lower maternal immunity against an emerging virus. As observed in this study, CV-A6 will likely affect more older children and present a similar age pattern to EV-A71 and CV-A16, since more maternal immunity is attained after introduction and spread of CV-A6, causing the median age of infection to go up. Further studies should be conducted to monitor the potentially changing epidemiology of CV-A6.

### Limitations

This study was subject to several limitations. First, due to unavailability of data, the number of HFMD associated hospitalizations in township hospitals was estimated from the proportion of HFMD-associated hospitalizations for all causes rather than from hospitalizations in pediatric or internal medicine departments. Nonetheless, the distribution of hospitalization causes is not expected to vary greatly since the demographics, and economic and living conditions were very similar among the different townships. Second, virological surveillance was not conducted at the 20 other township hospitals where the number of HFMD hospitalizations was very small (9%). Third, the HFMD patients hospitalized outside Anhua County were not captured. Due to low case-severity risk (1.1%) among HFMD patients, few cases need referral to prefecture- or provincial-level hospitals. Therefore, hospitalization rates of HFMD estimated in our study should be robust.

## Conclusions

Our study provides a comprehensive analysis of probable HFMD, lab-confirmed HFMD, EV-A71, CV-A16, CV-A6, CV-A10 and other enterovirus associated hospitalization rates in Anhua County, Hunan Province, China between 2013 and 2016. We found a substantial hospitalization burden annually for mild HFMD, caused by multiple enterovirus serotypes in China. During the study period, CV-A16 and CV-A6 contributed to more hospitalizations than EV-A71, CV-A10 and other enteroviruses. Our findings suggest that intensive training on how to quickly detect and treat mild HFMD should be provided to clinicians, especially those working at county-level and township-level hospitals. Furthermore, health education on HFMD should be provided to parents/guardians to address their concerns about HFMD.

## Acknowledgments

We thank staff members at the Anhua County-, Yiyang Prefecture-, and Hunan Provincial-level Departments of Health for providing assistance with administration and data collection; staff members at the Anhua County-, Yiyang Prefecture-, and Hunan Provincial-level CDCs and six study hospitals (Anhua People’s Hospital, Anhua Second People’s Hospital, Anhua Hospital of TCM, Tianzhuang Township Hospital, Jiangnan Township Hospital, and Qingtang Township Hospital) for providing assistance with field investigation, administration and data collection. We also thank Benjamin J. Cowling and Peng Wu for reviewing this manuscript (World Health Organization (WHO) Collaborating Centre for Infectious Disease Epidemiology and Control, School of Public Health, Li Ka Shing Faculty of Medicine, The University of Hong Kong, Hong Kong Special Administrative Region, China).

## Supporting information

**S1 Fig. Temporal trends of HFMD-associated hospitalizations by enterovirus serotype in six surveillance hospitals in Anhua County, China, October 2013 - September 2016.**

**S2 Fig. Annual hospitalization rates associated with HFMD stratified by age and enterovirus serotype in Anhua County, China, October 2013 - September 2016.** (A) Annual age-specific hospitalization rates of CV-A16-associated HFMD. (B) Annual age-specific hospitalization rates of CV-A6-associated HFMD. (C) Annual age-specific hospitalization rates of EV-A71-associated HFMD. (D) Annual age-specific hospitalization rates of CV-A10-associated HFMD.

**S1 Table. The age profile of HFMD-associated hospitalizations stratified by enterovirus serotype in 6 surveillance hospitals in Anhua County, China, October 2013 - September 2016.**

**S2 Table. Estimated hospitalization rates associated with HFMD by age group in Anhua County, China, October 2013 - September 2016.**

**S1 Checklist:** STROBE Checklist

